# The chromatin modifiers SET-25 and SET-32 are required for initiation but not long-term maintenance of transgenerational epigenetic inheritance

**DOI:** 10.1101/255646

**Authors:** Rachel Woodhouse, Gabrielle Buchmann, Matthew Hoe, Dylan Harney, Mark Larance, Peter R. Boag, Alyson Ashe

## Abstract

Some epigenetic modifications are inherited from one generation to the next, providing a potential mechanism for the inheritance of environmentally acquired traits. Transgenerational inheritance of RNA interference phenotypes in *Caenorhabditis elegans* provides an excellent model to study this phenomenon, and whilst studies have implicated both chromatin modifications and small RNA pathways in heritable silencing their relative contributions remain unclear. Here we demonstrate that the histone methyltransferases SET-25 and SET-32 are required for the establishment of a transgenerational silencing signal but not for long-term maintenance of this signal between subsequent generations, suggesting that transgenerational epigenetic inheritance is a multi-step process with distinct genetic requirements for establishment and maintenance of heritable silencing. Furthermore, small RNA sequencing reveals that the abundance of secondary siRNAs (thought to be the effector molecules of heritable silencing) does not correlate with silencing phenotypes. Together, our results suggest that the current mechanistic models of epigenetic inheritance are incomplete.

## Introduction

Despite the wide-held belief for the past hundred years that only information encoded in the genome of an organism can be inherited, it has recently become clear that epigenetic signals can sometimes be passed between generations (reviewed in Miska and Ferguson-Smith, 2016). Due to its short generation time and easily manipulated germline, *Caenorhabditis elegans* has emerged as one of the leading organisms with which to study this phenomenon, with a plethora of studies being reported over the last few years (Devanapally et al., 2015; Greer et al., 2011; Klosin et al., 2017; Shirayama et al., 2012).

One important tool that has been used to study transgenerational inheritance is RNAi silencing. RNAi was discovered in *C. elegans* (Fire et al., 1998) and from some of the first reports it was observed that occasionally the RNAi phenotype was detected in unexposed offspring for one or two generations (Fire et al., 1998; Grishok et al., 2000). Since then, several studies have shown inheritance of RNAi phenotypes for multiple generations (Alcazar et al., 2008; Ashe et al., 2012; Buckley et al., 2012; Gu et al., 2012; Houri-Ze’evi et al., 2016; Vastenhouw et al., 2006). In *C. elegans* RNAi is usually induced by feeding animals double stranded RNA (dsRNA) targeting the gene of interest. The double stranded RNA is processed by Dicer and accessory proteins to give primary small interfering RNAs (siRNAs) (Bernstein et al., 2001). 1° siRNAs have a 5’ monophosphate, are 21-23nt long, and are both sense and antisense to the target gene. The 1° siRNAs are bound by the argonaute protein RDE-1 (Tabara et al., 1999; Yigit et al., 2006) and are used to guide the production of secondary siRNAs. 2° siRNAs are almost exclusively 22 nucleotides long, predominantly have a 5′G, and a terminal triphosphate group (Pak and Fire, 2007; Sijen et al., 2001, 2007). 2° siRNAs (called 22Gs), which also exist in complex with an argonaute protein, are responsible for the degradation of the target mRNA in the cytoplasm (Aoki et al., 2007) or transcriptional gene silencing in the nucleus (Guang et al., 2010).

Genetic screens to find the components of the RNAi-initiated transgenerational epigenetic inheritance (TEI) pathway have implicated small RNA pathway components, especially the germline nuclear RNAi machinery, as well as histone modifying enzymes, implying that there may be some interplay between small RNAs and chromatin. During nuclear RNAi, the germline-specific nuclear Argonaute heritable RNAi defective 1 (HRDE-1) binds cytoplasmic 2° siRNAs and translocates to the nucleus (Ashe et al., 2012; Buckley et al., 2012; Shirayama et al., 2012). Upon interaction with complementary nascent mRNA transcripts, the nuclear RNAi defective (NRDE) factors NRDE-1, −2 and −4 are recruited (Ashe et al., 2012; Burton et al., 2011). The NRDE machinery then mediates gene silencing by inhibiting RNA polymerase II during transcriptional elongation (Guang et al., 2010), and by promoting chromatin modifications that are associated with gene silencing (Burkhart et al., 2011; Burton et al., 2011; Gu et al., 2012; Guang et al., 2010; Mao et al., 2015). HRDE-1 and NRDE-1, −2 and −4 are all required for TEI (Ashe et al., 2012; Buckley et al., 2012).

Repressive chromatin modifications such as H3K27me3 and H3K9me3 have been implicated in TEI in *C. elegans*. RNAi induces robust accumulation of H3K9me3, the hallmark of constitutive heterochromatin in most eukaryotes, at endogenous genes lasting for two generations (Gu et al., 2012; Kalinava et al., 2017). SET-25, an H3K9 methyltransferase (Snyder et al., 2016; Towbin et al., 2012), and SET-32, a putative H3K9 methyltransferase (Kalinava et al., 2017; Snyder et al., 2016; Spracklin et al., 2017), have both been implicated in TEI. Ashe et al. showed SET-25 is required for RNAi-initiated heritable silencing of a GFP transgene, and showed SET-32 is required in an analogous system triggered by PIWI-interacting small RNAs (piRNAs) (Ashe et al., 2012). Spracklin et al. have since shown that SET-32 is also required for RNAi-initiated heritable transgene silencing (Spracklin et al., 2017). SET-25, SET-32 and another H3K9 methyltransferase MET-2 are required for heritable accumulation of H3K9me3 in response to RNAi, but in contrast to TEI studies utilising transgene silencing, loss of SET-25 and SET-32 does not result in loss of heritable gene silencing of the endogenous *oma-1* gene (Kalinava et al., 2017), or promoter-mediated heritable germline silencing of the endogenous *sid-1* gene (Minkina and Hunter, 2017). Thus, there is still considerable debate as to the requirement of chromatin modifiers in TEI.

This study aims to determine the requirements for SET-25 and SET-32 in TEI. We use the same system as that used by Ashe and colleagues (Ashe et al., 2012), involving RNAi silencing of a germline-expressed GFP transgene. The visible nature of this phenotype provides an exquisitely sensitive system, whereby we can separate individual animals according to their silencing status and measure effects in these distinct groups. This approach has enabled us to probe the genetic requirements of TEI in each generation, and here we show that SET-25 and SET-32 are required for the establishment of a long-term silencing signal but not for its maintenance over subsequent generations, suggesting that TEI is a multi-step process. We also show that small RNAs are not as closely correlated with the presence of TEI as expected, and further characterise the phenotypes associated with mutations in *set-32* and *set-25*.

## Results

### set-25 and set-32 are required for transgenerational epigenetic inheritance to the F1 generation only

The requirement for *set-25* and *set-32* in multigenerational silencing in some studies led us to test whether they are required for transgenerational silencing of the germline-expressed *pie-1::gfp::h2b* transgene. We fed animals containing this transgene (‘sensor’) bacteria expressing anti-*gfp* dsRNA, triggering 100% silencing of the GFP transgene (P0 generation). In all strains, 0% of P0 control animals fed on empty vector bacteria were GFP-silenced (data not shown). Subsequent generations were produced by isolating silenced individuals, and were fed on regular OP50 bacteria. Selection of silenced individuals distinguishes this study from others that use RNAi transgene silencing models (Buckley et al., 2012; Burton et al., 2011; Houri-Ze’evi et al., 2016; Lev et al., 2017; Spracklin et al., 2017), and allows us to differentiate between effects in individual animals which inherit silencing and those which do not, and interrogate the requirements of genes at each generation. The percentage of GFP-silenced animals was measured at each generation. In wild-type sensor animals exposed to anti-*gfp* dsRNA, GFP silencing persisted for at least three generations after the dsRNA trigger was removed, as previously described (Ashe et al. 2012) (Figure 1A). Strikingly, *set-25(n5021)* and *set-32(ok1457)* F1 animals displayed a highly significant reduction in silencing proportions (Figure 1A). Surprisingly, the F2 and F3 offspring of animals that successfully inherited the silencing signal then displayed silencing proportions comparable to the sensor strain (Figure 1A). A CRISPR/Cas-9 generated mutant, *set-32(smb11)* (Figure S1), displayed the same inheritance pattern (Figure 1A). These data suggest that both *set-25* and *set-32* are required for transgenerational silencing in the F1 offspring, but are not required from the F2 generation onwards. The *set-32(ok1457); set-25(n5021)* double mutant shows the same inheritance pattern as *set-25(n5012)* alone, suggesting that *set-32* and *set-25* act in the same pathway (Figure 1A).

**Figure 1.**
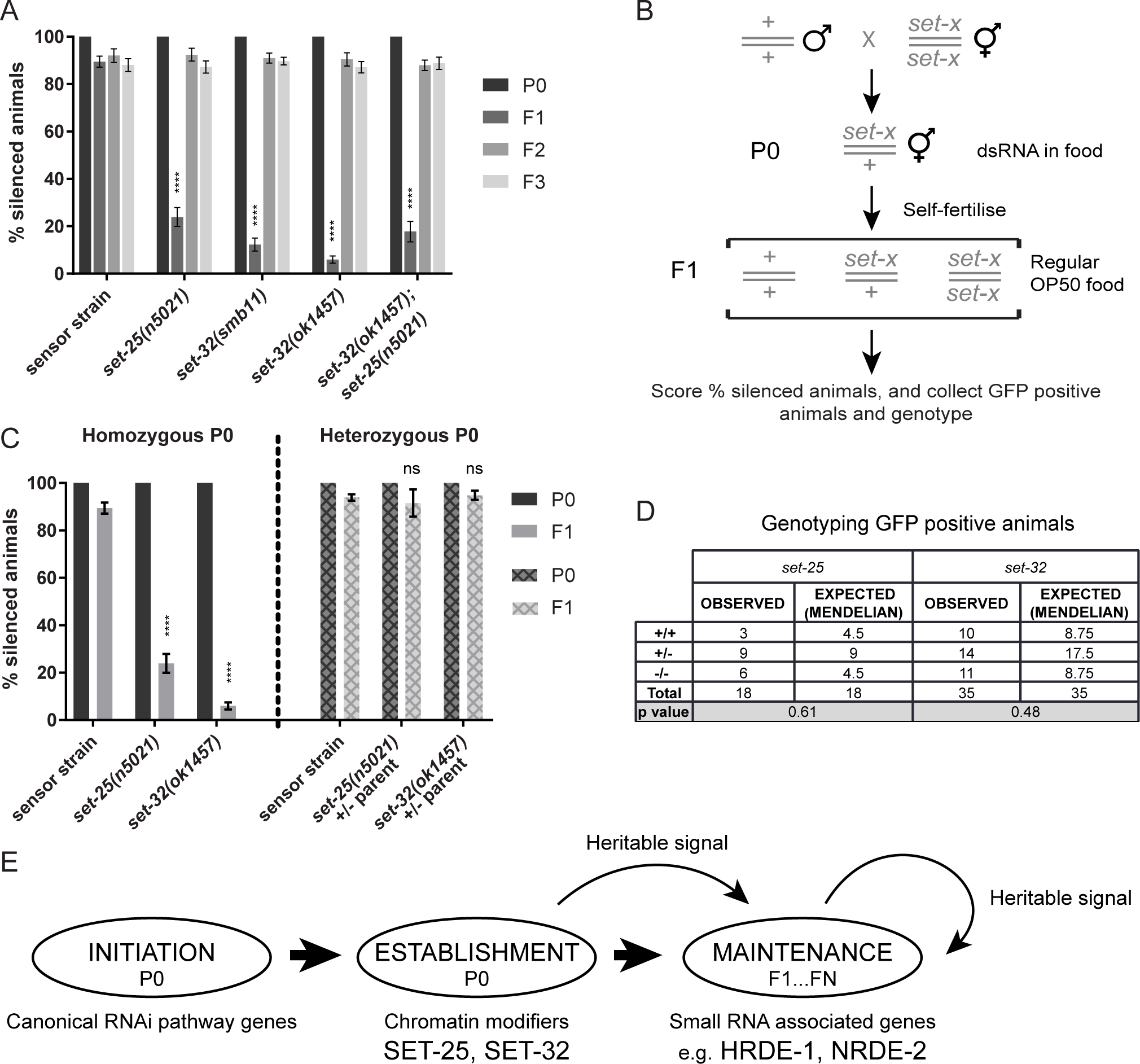
*set-25* and *set-32* are required for transgenerational epigenetic inheritance to the F1 generation. **A.** The percentage of GFP-silenced animals following exposure to GFP RNAi. P0 animals were exposed to GFP RNAi. F1-F3 animals were not exposed to the RNAi trigger, and were created by selecting silenced individuals and allowing them to self-reproduce. Data are mean ±SEM, n>100. Comparisons were performed by two-way ANOVA with Tukey’s post hoc. Statistics shown are mutant F1 compared to sensor strain F1, **** p<0.0001 **B.** Scheme of experiment to determine the requirement for *set-25* and *set-32* in the P0 or F1 generation for heritable silencing. *set-32* or *set-25* (*set-x*) heterozygous individuals were exposed to GFP RNAi. Their unexposed F1 progeny were scored for silencing inheritance, where an increased proportion of GFP positive worms compared with wild-type would indicate a requirement for SET-32 or SET-25 in the F1 generation. GFP positive worms were also collected and genotyped. **C.** The percentage of GFP-silenced animals produced by the assay in B. Data are mean ±SEM, n>100. Comparisons were performed by two-way ANOVA with Tukey’s post hoc. Statistics shown are mutant F1 compared to sensor strain F1, **** p<0.0001 **D.** The observed number of wild-type (+/+), heterozygous (+/−) and homozygous mutant (−/−) GFP positive F1 animals produced by the assay in B, and the expected number according to Mendel’s ratio. p values calculated by chi-square tests. **E.** Proposed three-step model of TEI.

Failure of silencing of the GFP transgene in the F1 offspring could be due to a requirement for *set-25* or *set-32* in either the P0 generation (*i.e.* helping to establish a heritable silencing signal) or the F1 generation (*i.e.* receiving or propagating a silencing signal). We sought to distinguish between these two possibilities by performing the silencing assay on *set-32* or *set-25* heterozygous individuals (P0) and assaying the inheritance of silencing in the F1 generation (Figure 1B). If *set-32* or *set-25* are required in the F1 generation we would expect the homozygous F1 mutants to display a failure of silencing (*i.e.* more homozygous mutants would express GFP than heterozygous or wild-type). Alternatively, if the proteins are required in the P0 generation only, the absence of functional protein in the F1 generation should not matter and we would expect to see wild-type levels of silencing amongst all offspring genotypes. Strikingly, we did not see an increased proportion of GFP-positive animals amongst the F1 homozygous mutants for either *set-25* or *set-32*; F1 offspring of *set-25* and *set-32* heterozygous parents displayed silencing proportions comparable to wild-type (Figure 1C), and homozygous mutants were not over-represented amongst GFP positive F1s (Figure 1D). This indicates that neither *set-25* nor *set-32* is required in the F1 generation, and therefore their role in epigenetic inheritance must be in the P0 generation. These results lead us to propose a three-step model of TEI consisting of initiation, establishment and maintenance phases, each requiring distinct factors (see Discussion) (Figure 1E).

### 22Gs do not correlate with heritable silencing

Since small RNA molecules, in particular 22Gs, have been implicated in heritable RNAi-induced silencing, we sequenced small RNAs in P0 animals and F1 animals in *set-25*, *set-32*, *hrde-1* and *nrde-2* mutant strains. P0 animals exhibited 100% silencing (Figure 1A) and so were collected in one pool, whilst F1 animals were separated into GFP-silenced (GFP-off) and GFP-expressing (GFP-on) pools, allowing us to investigate differences in small RNAs between these populations. Our hypothesis was that since all strains display GFP silencing in the P0 generation, they would have equal amounts of small RNAs mapping to the GFP transgene. Furthermore, we expected that F1 GFP-off worms would have anti-*gfp* small RNAs and that GFP-on worms would not. We first focussed our analysis on the wild-type animals. As expected, 1° siRNAs were present in the P0 generation but were essentially absent in the F1 generation (Figure 2Ai). 22Gs were present in the F1 GFP-off animals but were dramatically less abundant than in the P0 generation (Figure 2Aii,B), as expected (Ashe et al., 2012; Gu et al., 2012; Houri-Ze’evi et al., 2016). Surprisingly, 22Gs were also present in the GFP-on animals and the numbers of 22Gs did not differ significantly between GFP-off and GFP-on animals in wild-type (fold change off/on = 1.77, ns paired t-test) (Figure 2C,D). This is surprising because 22Gs are the effector molecules of silencing, and current models suggest that they are the molecule carried between generations by HRDE-1 (Minkina and Hunter, 2017); thus one would expect a substantial difference between GFP-off and GFP-on animals.

We next focussed on the various mutant strains. *set-25* animals have previously been shown to be defective in heritable siRNAs at the F3 generation (Lev et al., 2017). In our hands, we could detect GFP-targeting siRNAs in both the GFP-off and GFP-on F1 animals, at similar levels to those detected in wild-type (Figure 2A-D). Again, the difference in small RNAs between GFP-off and GFP-on animals was small. *set-32* and *hrde-1* animals showed slightly more anti-GFP 22Gs in the GFP-off animals compared to GFP-on animals (Figure 2C,D). Strikingly, the most abundant 22Gs were found in GFP-off *nrde-2* animals (Figure 2B-D), despite the defect in heritable silencing that these animals display. Why is there such an abundance of 22Gs in GFP-off animals in the *nrde-2* strain, which is defective in heritable silencing, when there is essentially no difference in 22G abundance between GFP-off and GFP-on animals in wild-type? It is tempting to speculate that 22Gs may not be the main heritable agent (see Discussion).

**Figure 2.**
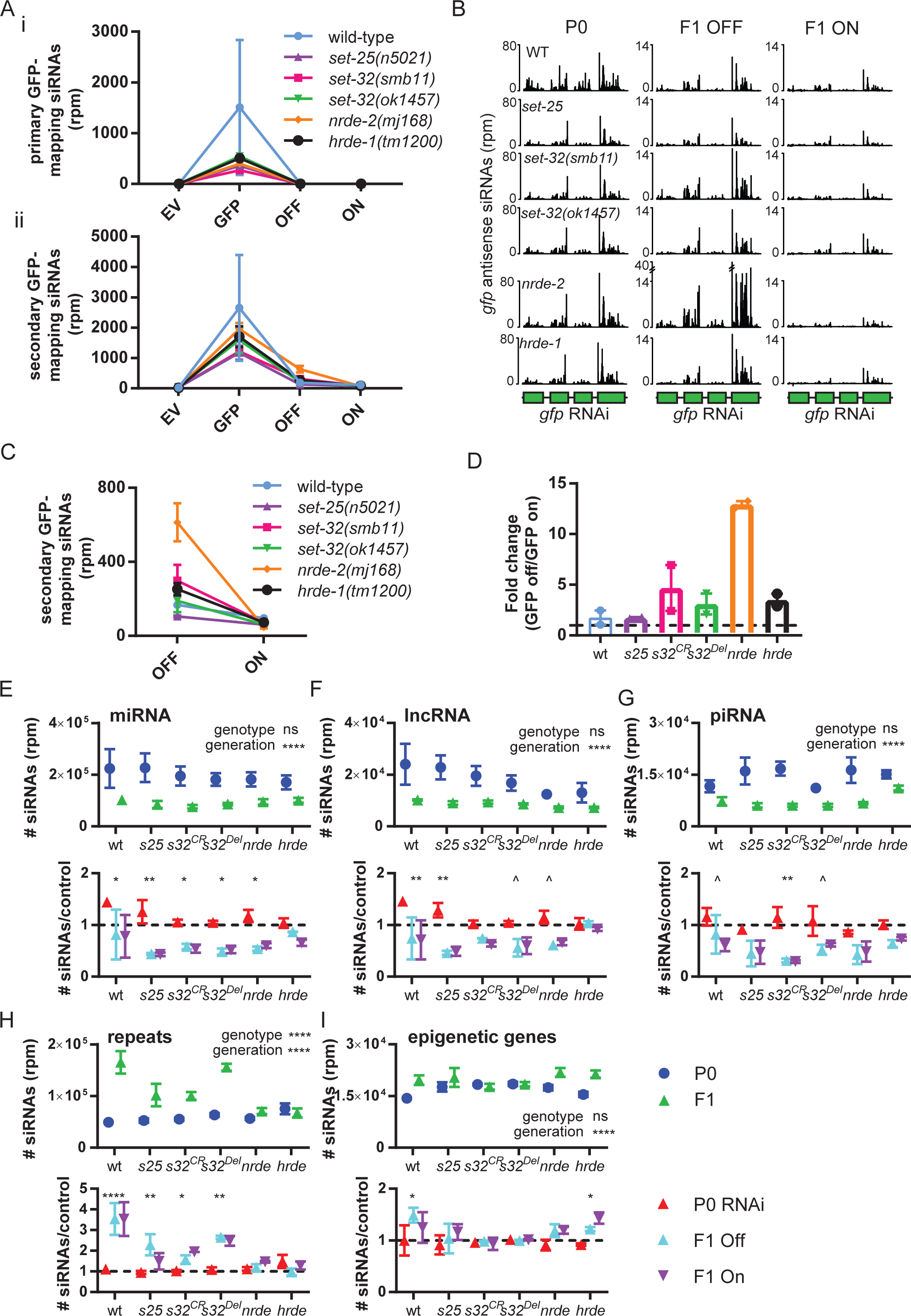
Small RNA analysis in the context of transgenerational epigenetic inheritance. **A.** The number of reads mapping to the GFP transgene in the sense (i) and antisense (ii) orientation for six different strains is shown. Points indicate the replicate while the line indicates the mean. Strains are identified by colour. **B.** Antisense *gfp*-mapping siRNAs (2° siRNAs) are shown for the P0 and F1 generations in various mutant strains as indicated. F1 animals were split into GFP silenced (‘OFF’) and GFP expressing (‘ON’) pools before library preparation. **C.** 2° siRNAs for the F1 generation only. **D.** Fold change of GFP mapping 2° siRNAs in GFP silenced (‘OFF’) vs GFP expressing (‘ON’) animals. Dashed line indicates equal expression between silenced and expressing animals. **E-I.** Small RNAs in reads per million (rpm) with the indicated biotype. Strains are indicated: wt is wild-type, *s25* is *set-25(n5021)*, *s32^CR^* is *set-32(smb11)*, *s32^Del^* is *set-32(ok1457)*, *nrde* is *nrde-2(mj168)*, *hrde* is *hrde-1(tm1200)*. The top panel shows P0 control and RNAi-treated combined (P0) and F1 GFP-expressing and GFP-silenced animals combined (F1). Two-way ANOVA was performed and results for variance at the genotype or generation level is shown. The bottom panel shows P0 and F1 individually, normalised to the relevant control (dashed black line). Comparisons were performed by two-way ANOVA with Tukey’s post hoc. ^ 0.05<p<0.1 * p<0.05 ** p<0.01 **** p<0.0001

### Endo siRNAs are perturbed in TEI mutants

It has previously been shown that RNAi can alter levels of endogenous siRNAs both in the exposed P0 generation and in the F1 generation in wild-type animals (Houri-Ze’evi et al., 2016; Lev et al., 2017). We first wanted to verify this finding. We took the small RNAs from control, P0 and F1 wild-type animals and divided them into seven classes of small RNAs based on the genomic annotation to which they map; transposable elements, protein coding, pseudogenic transcripts, repetitive regions, piRNAs, microRNAs (miRNAs) and long non-coding RNAs (lncRNAs). Contrary to previous reports (Houri-Ze’evi et al., 2016; Lev et al., 2017), we did not observe any differences in siRNA families between control and GFP RNAi exposed animals in any subset of small RNAs in wild-type animals (Figure S2A). We also did not observe any differences between F1 GFP-off and GFP-on animals. We performed the same analysis in the *set-25*, *set-32*, *nrde-2*, and *hrde-1* mutant strains and also saw no difference between control or RNAi-treated animals in the P0 generation, or between GFP-off and GFP-on animals in the F1 generation.

We did, however, observe some significant differences between the P0 and F1 generations. miRNA-mapping siRNAs were decreased in the F1 generation compared to the P0 generation (Figure 2E), in direct contrast to previous reports (Houri-Ze’evi et al., 2016) where an increase in miRNAs was observed in the F1 generation. lnc-RNA-mapping siRNAs also displayed a decrease in F1 (Figure 2F). These patterns were observed in wild-type and most mutant strains (Figure 2E,F), suggesting that there may be a heritable response elicited by RNAi exposure in wild-type animals that decreases the levels of these classes of endo siRNAs in the F1 generation, perhaps due to competition for components of the RNAi machinery. *hrde-1* animals had no difference between P0 and F1 generations, suggesting that in this strain the heritable endo siRNAs pathways are perturbed. A lower amount of endo siRNAs in F1 compared to P0 was detected over all strains for piRNAs (further enhanced in *set-32* mutants) (Figure 2G), transposable elements and pseudogenic-mapping siRNAs (Figure S2). *nrde-2* animals had globally higher levels of pseudogenic-mapping endo siRNAs in both generations compared to wild-type.

Repeat-mapping siRNAs increased greatly in the F1 generation compared to the P0 generation in wild-type animals (Figure 2H). This increase was lost in all mutant strains except *set-32(smb11*), again suggesting that a heritable response to RNAi exposure is perturbed in these mutant strains, consistent with our previous results. A role for *set-25* and *nrde-2* in the silencing of repetitive elements has recently been reported (McMurchy et al., 2017). It is interesting that we do not detect an overall decrease in siRNA mapping in these strains, and only see the effect in the F1 generation. Perhaps a subtle effect in the mutant strains is exacerbated by competition with the heritable RNA response.

Finally, we looked at siRNAs that map to protein coding genes. We saw a slight increase in these endo siRNAs in *nrde-2* compared to wild-type but otherwise found no obvious differences (Figure S2). However when we took a subset of genes – those that are involved in epigenetic processes (Houri-Ze’evi et al., 2016) – we saw that siRNAs mapping to those genes were upregulated in the F1 generation in wild-type, but that this effect was lost in all strains except *hrde-1* (Figure 2I).

Overall, we saw that endo siRNAs alter following RNAi exposure in the F1 generation in wild-type animals, and that this heritable endo RNAi response was changed in all of the TEI mutant strains that we analysed. Our data suggest that heritable siRNA pathways are disturbed in these strains globally and not only at the GFP locus that we targeted in the RNAi.

### SET-25 and SET-32 have distinct germline expression patterns

In order to further characterise SET-32 and SET-25 we used CRISPR/Cas9 to insert *mCherry* immediately downstream of the start codon of *set-25* and immediately upstream of the stop codon of *set-32* to generate N‐ and C-terminal tagged proteins respectively. These strains displayed wild-type heritable silencing in the RNAi inheritance assay, indicating that the tagged proteins are functional (data not shown). For mCherry::SET-25 we saw expression in the nuclei of the mitotic zone of the germline. This expression was detected from larval stage 2 (L2) onwards (Figure 3A,B and data not shown). Faint expression was also detected in all nuclei of embryos from about the 8-16 cell stage. Although expression was weak, mCherry::SET-25 was seen to be associated with the chromatin throughout mitotic cell divisions in the embryos (Figure 3C).

**Figure 3.**
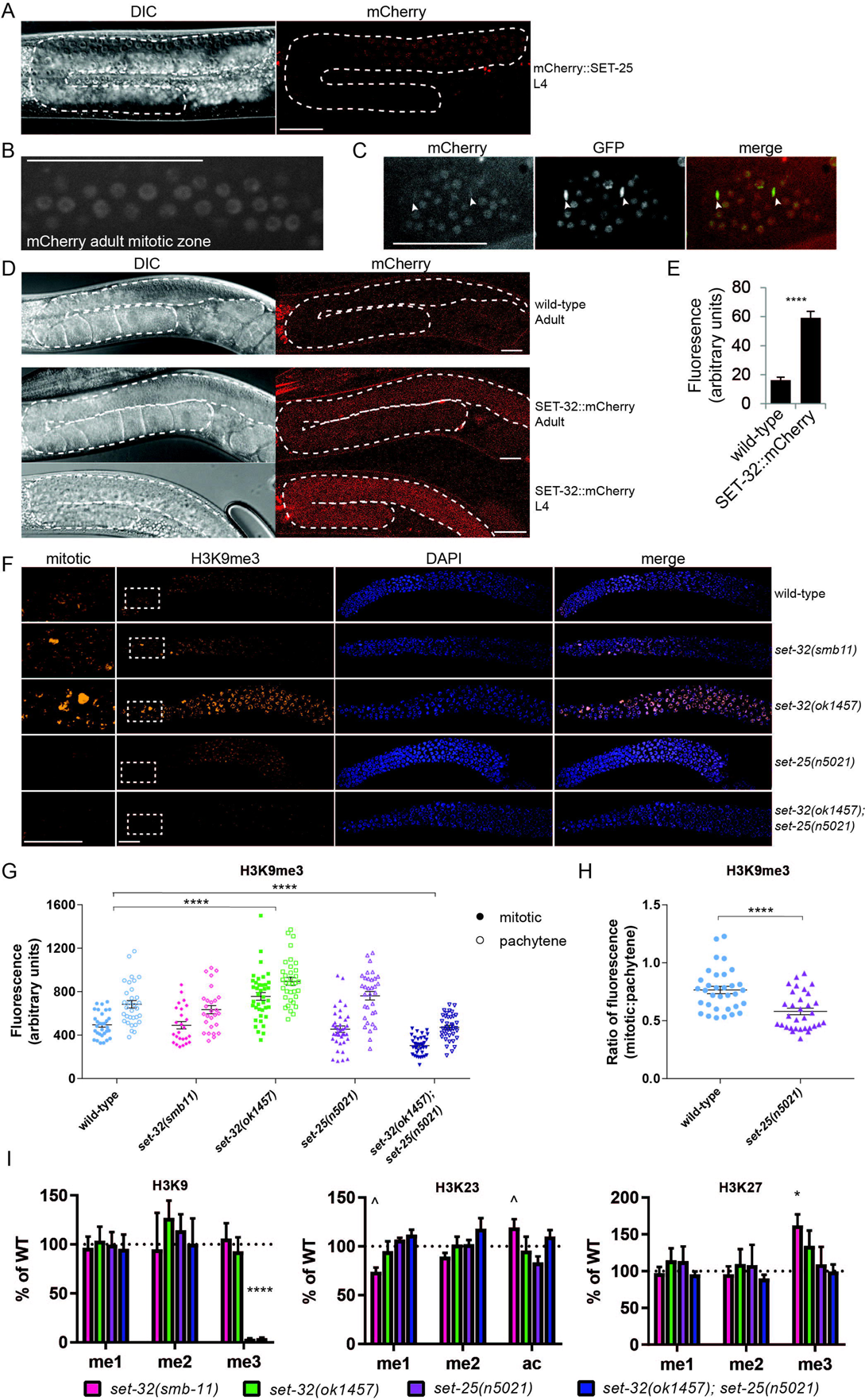
SET-32 and SET-25 expression patterns and H3K9 methylation analysis. **A.** One germline of a representative L4 mCherry::SET-25 expressing animal. Germline is outlined in white. **B.** Nuclear localisation of mCherry::SET-25 in the mitotic zone of an adult *C. elegans*. **C.** Embryo expressing mCherry::SET-25 (left). Chromatin is marked by HIS-58::GFP (middle). Arrows indicate dividing cells with condensed chromatin. **D.** The entire germline of a wild-type adult (top), and SET-32::mCherry expressing adult (middle) and L4 (bottom) animals. Germline is outlined in white. **E.** Fluorescence intensity in the entire germline of SET-32::mCherry (n=15) and wild-type (n=11) animals was quantified using ImageJ. Comparison was performed by t-test. **** p<0.0001. **F.** Representative single confocal plane images of dissected one-day-old adult gonads of the indicated strain. The mitotic zone is on the left and the pachytene zone is on the right. Panels show a portion of the mitotic zone (enlarged from the region indicated by the dashed white box), anti-H3K9me3 staining (yellow), DAPI staining (blue), and an overlay of H3K9me3 and DAPI staining, as indicated. Strains were imaged under identical parameters within staining conditions. All scale bars in **A-F** represent20 µm. **G.** Quantification of fluorescence intensity of H3K9me3 staining in the mitotic and pachytene zones. ImageJ was used to generate an intensity threshold-based mask for nuclei using the DAPI signal, then the average intensity of H3K9me3 staining in nuclei was measured. Data are mean ±SEM, n=25-39. Comparisons were performed by two-way ANOVA with Tukey’s post hoc. **H.** Ratio of fluorescence intensity in mitotic:pachytene zones. Data are mean ±SEM, n=25-39. Comparison was performed by t-test. **** p<0.0001 **I.** Quantification of H3K9, H3K23 and H3K27 levels in the indicated mutant strains by mass spectrometry. For each modification, levels were normalised to a relevant control peptide and are displayed as % of wild-type (indicated by dotted line). Data are mean ±SEM, n=3 replicates for each mutant strain. Comparisons were performed by two-way ANOVA with Dunnett’s post hoc. ^ p=0.058, * p<0.05, **** p<0.0001

Very weak SET-32::mCherry expression was also observed in the germline, but was not localised exclusively to nuclei, nor to the mitotic zone. Instead, expression was detected throughout the germline from L1/2 onwards, with maximum expression detected at L4 (Figure 3D and Figure S3A). No expression could be detected in embryos. As the expression of SET-32::mCherry was so weak, we imaged SET-32::mCherry and wild-type animals under identical conditions and quantified the amount of fluorescence in whole adult germlines using ImageJ. There was significantly more fluorescent signal detected in SET-32::mCherry animals compared to wild-type (Figure 3E). Cytoplasmic expression of a histone methyltransferase is not without precedence, as a previous study showed the H3K9 methyltransferase MET-2 to be enriched in the cytoplasm (Towbin et al., 2012). It is worth noting that SET-32::mCherry does not appear to be excluded from the nucleus, implying that it may have both nuclear and cytoplasmic roles. mCherry::SET-32 was not detectable by Western blot (data not shown), thus we cannot rule out the possibility that the mCherry tag is cleaved from the SET-32 protein. Despite this caveat, our data show that *set-32* is expressed exclusively in the germline.

### set-25 and set-32 mutants have perturbed germline H3K9me3

Since SET-25 and SET-32 are both expressed in the germline and have been implicated in H3K9 methylation (Kalinava et al., 2017; Snyder et al., 2016; Spracklin et al., 2017; Towbin et al., 2012), we performed immunofluorescence against H3K9me3 and H3K9me2 in dissected gonads of mutant hermaphrodite adults. We observed a significant reduction in H3K9me3 staining in *set-32(ok1457); set-25(n5021)* mutants throughout the germline compared with wild-type (Figure 3F,G) and a corresponding increase in H3K9me2 staining (Figure S3B,C). No other strains displayed a difference in H3K9me2 staining (Figure S3B,C).

Surprisingly, no difference was observed in H3K9me3 intensity in the mitotic or pachytene zones of *set-25(n5021)* or *set-32(smb11)* germlines compared with wild-type (Figure 3F,G). However, in *set-25(n5021)* mutant germlines we observed a significant decrease in intensity in the mitotic region relative to the pachytene region compared with wild-type (Figure 3H), consistent with a role for SET-25 function in the mitotic region. Both of these alleles are putative null mutants with predicted loss of the SET domain responsible for addition of methyl groups to histones; *set-25(n5-21)* carries a 1979 base pair deletion which removes over half of the SET domain coding sequence, and *set-32(smb-11)* carries a 32 base pair frameshift mutation in exon 2 resulting in a predicted premature stop codon before the SET domain (Figure S1). Thus, the lack of large differences in H3K9me3 in these putative null mutant strains implies a level of redundancy of H3K9 trimethyltransferases in the germline (in contrast to the role of SET-25 previously shown in embryos (Towbin et al., 2012)), and/or that the role of SET-25 and SET-32 in H3K9me3 deposition is at specific loci or developmental stages, or in response to particular stimuli and therefore not detected in this experiment. The latter model is consistent with the results of Spracklin et al. who recently showed that loss of SET-32 results in decreased H3K9me3 specifically at genes targeted by HRDE-1 and not at non-HRDE-1 target genes (Spracklin et al., 2017).

Interestingly the *set-32(ok1457)* allele behaved differently to the *set-32(smb11)* allele; H3K9me3 intensity in *set-32(ok1457)* mutants was significantly higher than wild-type (Figure 3F,G) and highly variable between nuclei (Figure 3F, Figure S3D). Variation in H3K9me3 has also been observed between *set-32(ok1457)* intestinal nuclei (Snyder et al., 2016) (although this is inconsistent with our observation of SET-32 expression exclusively in the germline). This variation could be explained by the *set-32(ok1457)* allele; it carries a 514 base pair in frame deletion which removes most of exons 2 and 3 but potentially leaves the SET domain intact and functional (Figure S1). The deletion may remove important sequences required for correct targeting of the protein, resulting in misregulation and hence aberrant methyltransferase activity, manifesting as variable germline H3K9me3. It is interesting that we do not observe this aberrant H3K9me3 in the double mutant strain, which contains the *set-32(ok1457)* allele but lacks SET-25. This suggests that *set-32(ok1457)* requires the presence of a functional SET-25 protein to exert its aberrant effect.

Strikingly, all mutant strains including the double mutant displayed some level of H3K9me3 in the germline (Figure 3F), suggesting that there is at least one other H3K9 trimethyltransferase acting in the germline or that the antibodies bind non-specifically at low levels. One possible H3K9 trimethyltransferase candidate is SET-26, which has previously been implicated in H3K9me3 deposition by *in vitro* methyltransferase assay (Greer et al., 2014).

### SET-32 does not affect H3K9 methylation in whole worms

Some reports have linked SET-32 to H3K9me3 accumulation, but each showed a correlation between a loss of SET-32 and decrease in H3K9me3 (Kalinava et al., 2017; Snyder et al., 2016; Spracklin et al., 2017). Since we observed no decrease in H3K9me3 in *set-32* mutant germlines by immunofluorescence, we performed bottom-up proteomics on histones extracted from wild-type and *set-32* mutant strains as an unbiased approach to identifying the modification(s) for which SET-32 is responsible.

While we could see clear evidence for the role of SET-25 in trimethylation of H3K9 (as previously shown by Towbin and colleagues (Towbin et al., 2012)), we could see no evidence for SET-32 playing a similar role (Figure 3I). Our analysis was performed on whole animals from mixed-stage populations. Given the germline localisation of SET-32 determined by our mCherry expression strain and low expression levels, it is possible that a role of SET-32 in H3K9 methylation at specific loci and/or developmental times is obscured.

We detected a trend towards a decrease in H3K23 mono-methylation in *set-32(smb11*) mutants and a corresponding increase in H3K23 acetylation (Figure 3I). These changes were not detected in the *set-32(ok1457)* allele. We also detected an increase in H3K27 tri-methylation in *set-32(smb11)* (Figure 3I). It is unlikely that a putative methyltransferase mutant would directly cause an increase in methylation, so this is most likely an indirect effect indicating a general disruption of histone modifications in *set-32* mutants. A similar trend was observed in the *set-32(ok1457)* strain. We detected a decrease in H3K79 di-methylation in the *set-32(ok1457); set-25(n5021)* strain, with a similar trend evident in both single mutant strains (Figure S3E). It has been suggested that H3K79 methylation occurs in a nucleosomal context and is regulated by cross-talk between histone modifications (Farooq et al., 2016), so this additive effect may be an indirect result of changes to the overall histone modification pattern in the double mutant.

### set-25 mutants do not display a mortal germline phenotype

A recent report (Spracklin et al., 2017) showed that *set-32* mutants display a mortal germline (Mrt) phenotype – that is, after just two generations at 25°C the brood size dropped by 75%, and by 3-5 generations the animals became sterile. We were interested in testing if this was also the case for *set-25* mutants. We shifted *set-32(ok1457)*, *set-32(smb11)* and *set-25(n5021)* mutants to 25°C and serially passaged individual animals for 12 generations, counting brood size at each generation. We observed a Mrt phenotype in *set-32* mutants, although it was milder than previously reported (Spracklin et al., 2017); even after twelve generations at 25°C, *set-32(ok1457)* and *set-32(smb11)* were not fully sterile (Figure 4A). Surprisingly however, we did not observe a Mrt phenotype in *set-25* mutants (Figure 4A). We continued following the brood size of the wild-type and *set-25* mutant lines and found no difference between them at the 20^th^ and 25^th^ generation (data not shown). We also counted the number of sterile individuals which arose during maintenance at 25°C for 12 generations, and again observed no difference between *set-25* mutants and wild-type (Figure S4A).

**Figure 4.**
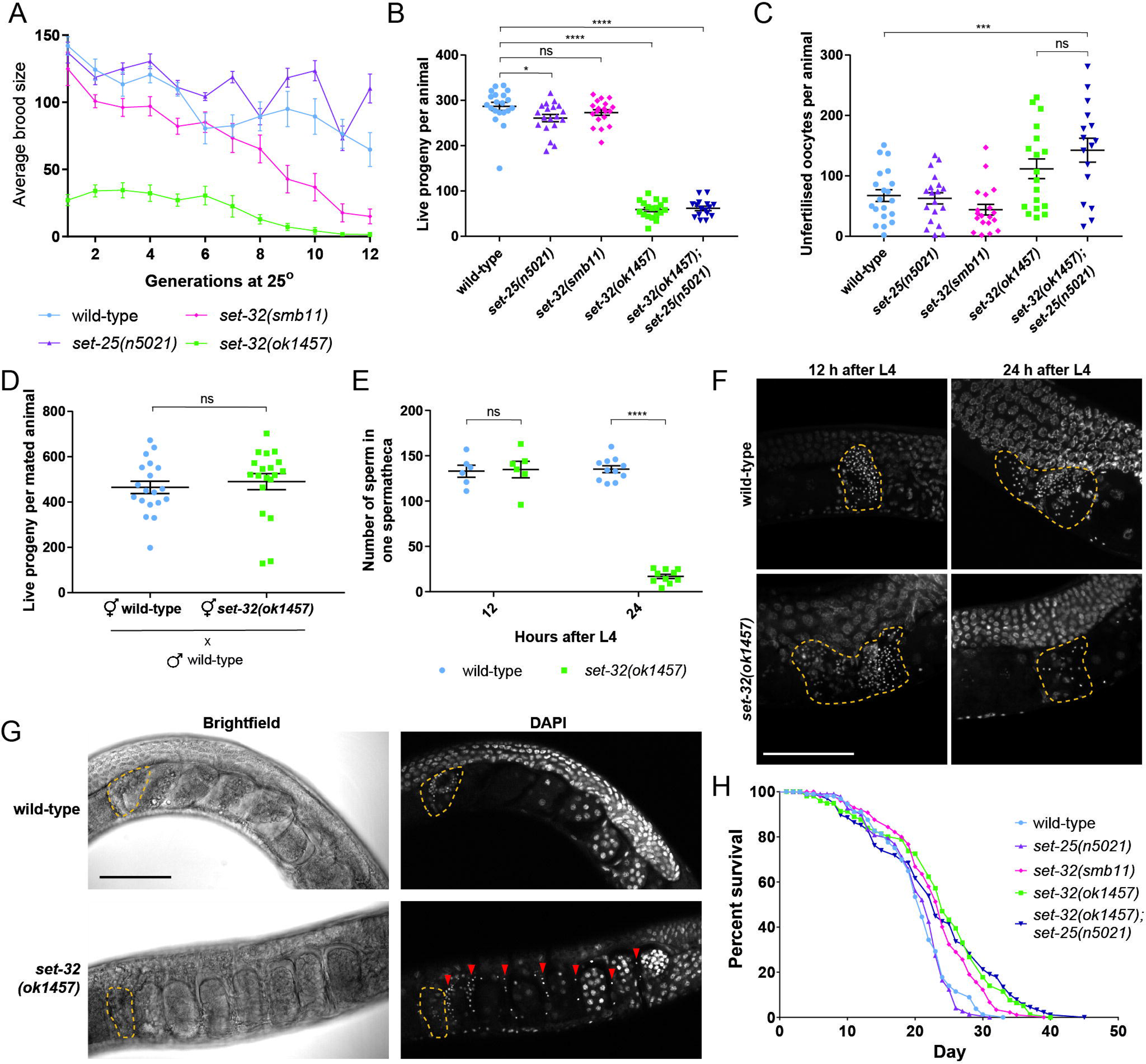
*set-32* mutants display a mortal germline phenotype, defective sperm, and extended lifespan. **A.** Animals were shifted to 25°C at L3 and the brood size of successive generations created by passaging individual worms was counted. Data are mean ±SEM, n=20 lines. **B-C.** The total number of **B.** live progeny and **C.** unfertilised eggs per animal was counted. Data are mean ±SEM, n=18-20. Comparisons were performed by one-way ANOVA with Dunnet’s post hoc and Tukey’s post hoc respectively. * p<0.05 *** p<0.001 **** p<0.0001 **D.** Brood sizes of wild-type or *set-32(ok1457)* hermaphrodites mated to wild-type males were counted. Data are mean ±SEM, n=19. Comparison was performed by t-test. **E.** Sperm nuclei were visualised by DAPI and the number of sperm in one spermatheca per animal was counted 12 hours (n=6) and 24 hours (n=10-12) after L4. Data are mean ±SEM. Comparisons were performed by two-way ANOVA with Sidak’s post hoc. **** p<0.0001 **F.** Representative maximum intensity Z-projection of a spermatheca in DAPI-stained animals, 12 hours and 24 hours after L4. **G.** Representative images of DAPI-stained animals 24 hours after L4. In *set-32(ok1457)* animals, sperm can be detected throughout the uterus, indicated by red arrowheads. In **F-G** scale bars represent 50 µm and the position of the spermatheca is indicated by dashed yellow line. **H.** Survival curves for animals of the indicated genotype in the absence of FUDR. p<0.0001 for *set-32(smb11)* vs. wild-type and *set-32(ok1457)* vs. wild-type, ns for *set-25 (n5021)* vs. wild-type. n=110 at Day 0, comparisons were performed by Log-rank test.

### set-32(ok1457) animals have defective sperm

As SET-32 mutants have not been widely studied, we were interested in further characterising our mutant strains. We noticed that *set-32(ok1457)* animals did not starve plates as quickly as wild-type animals grown under the same conditions and investigated this further by performing brood size assays, counting the numbers of live progeny and unfertilised eggs. *set-32(ok1457)* animals displayed a dramatically reduced brood size compared with wild-type, producing 59 and 287 live self-progeny on average respectively (Figure 4B). *set-25(n5021)* animals displayed a small but statistically significant reduction in brood size compared with wild-type (mean of 287 vs 261 progeny). *set-32(ok1457); set-25(n5021)* double mutants (mean of 62 progeny) displayed the same phenotype as *set-32(ok1457)* alone. Interestingly, the putative null mutant *set-32(smb11)* displayed no brood size defects, implying that the defects exhibited by *set-32(ok1457)* are due to the nature of the allele. As proposed in the immunofluorescence results, protein expressed from the *set-32(ok1457)* allele may be misregulated. This deleterious gain of function could be responsible for the observed brood size defects.

Production of live progeny from wild-type and *set-25(n5021)* animals peaked at day 2 of adulthood, and animals continued to lay live progeny until day 4 (Figure S4B). In contrast, *set-32(ok1457)* and *set-32(ok1457); set-25(n5021)* live progeny production peaked at day 1 of adulthood, with animals laying very few live progeny from day 2 onwards (Figure S4B). In both these strains animals did not cease to lay eggs at day 1, but instead continued laying a large number of unfertilised oocytes (Figure S4C). In total, *set-32(ok1457); set-25(n5021)* animals produced significantly more unfertilised oocytes than wild-type (mean of 143 vs 68) (Figure 4C). *set-32(ok1457)* animals also displayed a strong trend of increased production of unfertilised oocytes (mean of 112), that was not significantly different from the number of *set-32(ok1457); set-25(n5021)* unfertilised oocytes (Figure 4C). Together, these data strongly suggest that *set-32(ok1457)* animals lay more unfertilised oocytes than wild-type, and that this is not greatly enhanced in the double mutant.

The large numbers of unfertilised eggs suggested a sperm defect. In order to test this hypothesis, we crossed wild-type male animals with *set-32(ok1457)* hermaphrodites. We observed that the brood size from *set-32(ok1457*) females mated with wild-type males was indistinguishable from wild-type mated brood size (Figure 4D, Figure S4D), suggesting that *set-32(ok1457)* hermaphrodites have male germline defects only.

In order to further characterise the male germline defects, we performed DAPI staining on wild-type and *set-32(ok1457)* hermaphrodites 12 and 24 hours after the L4 stage, and counted the number of sperm nuclei. At 12 hours post L4, there was no difference in the number of sperm per spermatheca between the two strains (wild-type = 133 sperm, *set-32(ok1457)* = 135 sperm) (Figure 4E,F). However, at 24 hours post L4 there was a striking, highly significant reduction in the number of sperm in the *set-32(ok1457)* spermathecas (wild-type = 135 sperm, *set-32(ok1457)* = 17 sperm) (Figure 4E,F). Furthermore, at 24 hours post L4 we detected large numbers of sperm throughout the uterus of *set-32(ok1457)* animals which were not present in wild-type animals (Figure 4G). In wild-type animals some sperm are pushed from the spermatheca into the uterus by the passage of fertilised oocytes but quickly migrate back, resuming positions in the spermatheca before the passage of the next oocyte (Ward and Carrel, 1979). Thus, the presence of large numbers of sperm in the uterus suggests that *set-32(ok1457)* self-sperm are motility-defective.

We wanted to test whether *set-32(ok1457);* male sperm are also defective, so we crossed *set-32(ok1457); him-8(e1489)* males or control *him-8(e1489)* males, both carrying a neuronal GFP transgene, to wild-type hermaphrodites. The *him-8* mutation produces high incidence of males and was used to aid in obtaining large quantities of males. Cross-progeny express the GFP transgene whilst self-progeny do not, allowing us to distinguish between the two. Male sperm typically outcompete hermaphrodite self-sperm (Ward and Carrel, 1979), and we observed that wild-type mated controls produced predominantly cross-progeny as expected (Figure S4E). In contrast, wild-type hermaphrodites mated to *set-32(ok1457)* males produced significantly less cross-progeny (Figure S4E), indicating a male sperm defect. Cross‐ and self-progeny were produced in parallel throughout the first few days of adulthood (data not shown) suggesting that mutant male sperm fail to outcompete wild-type hermaphrodite self-sperm, possibly due to a motility defect as observed in mutant hermaphrodite self-sperm.

### Loss of SET-32 extends lifespan

We were also interested to see whether loss of SET-25 or SET-32 altered lifespan. We performed lifespan assays in the absence of FUDR on wild-type, *set-25(n5021), set-32(ok1457)* and *set-32(smb11)* mutants. In data representative of two independent assays, *set-25(n5021)* lifespan was not significantly different to wild-type (median lifespan of 22 vs 21 days) (Figure 4H). However, both *set-32(ok1457)* and *set-32(smb11)* animals displayed significant lifespan extension compared with wild-type (median lifespan 24 vs 21 days, 14% increase) (Figure 4H, Figure S4F,J). There is a well-known inverse relationship between fertility and lifespan (Partridge et al., 2005). The fact that we see a lifespan extension in both *set-32* mutant strains and reduction of fertility in only one argues that reduced fertility cannot be the cause of the lifespan extension observed here. Nonetheless, in order to rule out this possibility we also performed the lifespan assays in the presence of FUDR which induces sterility. Again, we saw an extension of lifespan in *set-32(ok1457)* animals compared to wild-type (median lifespan of 28 vs 24, 17% increase) (Figure S4G-J), indicating lifespan extension is independent of reduced brood size.

## Discussion

### SET-25 and SET-32 establish a heritable silencing signal

Here we have shown that the histone methyltransferases SET-25 and SET-32 are required only in the generation exposed to the initial silencing trigger to establish silencing in the next generation, and that when established, transgenerational silencing can be efficiently propagated in all subsequent generations in the absence of SET-32 and SET-25. In stark contrast, HRDE-1 and NRDE −2 are required to maintain heritable silencing in subsequent generations (Ashe et al., 2012; Buckley et al., 2012; Minkina and Hunter, 2017). Even when silenced animals are selected to create the next generations, *hrde-1* and *nrde-2* mutants display heritable silencing failure in increasing proportions in successive generations (Ashe et al., 2012). None of these genes are required to silence the generation initially exposed to the RNAi trigger (Ashe et al., 2012; Buckley et al., 2012; Spracklin et al., 2017, this study). Thus, we propose a three-step model of TEI consisting of initiation of silencing by canonical RNAi pathway genes, establishment of heritable silencing by *set-25* and *set-32*, and ongoing maintenance of heritable silencing requiring small RNA-associated genes such as *hrde-1* and *nrde-2* (Figure 1E). Further work will need to be performed to place other recently identified TEI genes correctly into this model (Akay et al., 2017; Houri-Ze’evi et al., 2016; Spracklin et al., 2017; Weiser et al., 2017). This model is highly complementary to the results of Kalinava et al. (“The establishment of nuclear RNAi is a transgenerational process and is promoted by a putative H3K9 methyltransferase SET-32 in *Caenorhabditis elegans”*, co-submitted).

How might this three-step model work? As SET-25 and SET-32 are histone methyltransferases, it follows that the establishment of a heritable silencing signal involves the deposition of H3K9me3 at the targeted locus. Indeed, SET-32 is required for accumulation of H3K9me3 at loci targeted by both exogenous and endogenous siRNAs (Kalinava et al., 2017; Spracklin et al., 2017). However, previous studies have demonstrated that silencing can be inherited across generations in the absence of the targeted DNA locus (Grishok et al., 2000; Houri-Ze’evi et al., 2016; Minkina and Hunter, 2017; Rechavi et al., 2011; Sapetschnig et al., 2015), suggesting that H3K9me3 cannot be the signal inherited between generations. In each of these experiments at least one copy of the target locus is present in animals exposed to the silencing trigger. We propose that the heritable silencing signal established by SET-25 and SET-32 is established in the mitotic zone of the germline of exposed animals when at least one copy of the targeted locus is present in every cell, triggering a locus-independent mechanism that then maintains heritable silencing throughout the subsequent zones of the germline and into the inheriting generations. This is consistent with the expression of SET-25 in the mitotic zone of the germline.

What could this locus-independent mechanism be? In its role as a H3K9 methyltransferase, SET-25 is required for complete anchoring of heterochromatic arrays to the nuclear periphery (Towbin et al., 2012). Furthermore, SET-25 co-localises with its own methylation product in perinuclear foci, suggesting that following anchoring it mediates propagation of heterochromatin to neighbouring loci (Towbin et al., 2012), consistent with observations of the spread of H3K9me3 to loci neighbouring RNAi targeted genes (Burton et al., 2011; Gu et al., 2012). Potentially then, the locus-independent silencing mechanism could involve localisation of the silenced region to the nuclear periphery, a region associated with gene silencing (Meister and Taddei, 2013). In this model, SET-25 and SET-32 could deposit H3K9me3 at the target locus, resulting in the anchoring of this locus to the nuclear periphery. Subsequently, SET-25 and SET-32 could mediate the propagation of heterochromatin to neighbouring loci, including to homologous regions adjacent to an absent locus on the other chromosome, tethering the local region to the nuclear periphery and establishing a silencing signal which is independent of the specific locus. SET-32 may act to maintain the position of these loci at the nuclear periphery during meiosis, since SET-32 expression was observed throughout the germline.

Silencing in subsequent generations is maintained by HRDE-1 and NRDE-2, and 22Gs. However, our results suggest that siRNAs cannot be the main heritable agent, because abundance of 2° siRNAs did not correlate with heritable silencing in wild-type or mutant strains. Potentially, 2° siRNAs mediate heritable silencing in conjunction with another mechanism, which may involve maintenance of the silenced locus at the nuclear periphery, or a different type of RNA molecule. Future research should investigate the nuclear localisation of the targeted locus during heritable RNAi to determine whether it is anchored to the nuclear periphery, and investigate how nuclear localisation might interact with RNAs and associated factors to maintain heritable silencing across generations.

### Chromatin modifications mediated by SET-32

Several reports have linked SET-32 to H3K9me3 accumulation (Kalinava et al., 2017; Snyder et al., 2016; Spracklin et al., 2017). We have investigated the chromatin modifications for which SET-32 is responsible using two approaches. Using bottom up proteomics we have shown that *set-32* mutants display no global decrease in H3K9me3 compared with wild-type in whole worms. We did, however, see evidence for a role for SET-32 in directly or indirectly mediating several other modifications, including H3K23me, H3K23ac, H3K27me3, and H3K79me2. Using immunofluorescence we have shown that *set-25* and *set-32* putative null mutants exhibit germline H3K9me3 comparable to wild-type, although a significant decrease was observed in a double mutant. Furthermore, the potentially misregulated *set-32(ok1457)* mutant showed increased germline H3K9me3. These latter results further support the hypothesis that SET-32 mediates H3K9me3 accumulation. However, it is clear that its role is complex; the lack of H3K9me3 loss at the whole germline and whole worm level in *set-32* mutants suggests that SET-32 may be functioning at least partially redundantly with SET-25 in H3K9me3 accumulation in the germline, but the presence of severe defects in heritable silencing, germline maintenance, and lifespan regulation in the single mutant implies that it must have non-redundant function in these processes. Future work should address which modification(s) SET-32 is directly modifying and which are indirectly affected, and whether SET-32 operates in conjunction with other methyltransferases to carry out its methyltransferase activity.

### The role of the RNAi inheritance machinery in maintaining germline immortality and implications for lifespan

In *C. elegans* several genes found to be involved in TEI have also been reported to have Mrt phenotypes, leading to the hypothesis that germline immortality maintenance is a general role of the RNAi inheritance machinery (Buckley et al., 2012; Spracklin et al., 2017). In *set-25* we show the first example, to our knowledge, of a gene required in TEI which does not exhibit a Mrt phenotype; after 25 generations at 25°C, *set-25* mutants displayed comparable fertility to wild-type animals. Our results suggest two potential explanations; *set-25* mutants exhibit a mild Mrt phenotype which may take many generations to appear and hence was not detected in our assay, or SET-25 is not required for maintaining germline immortality. It is interesting that TEI mutants display Mrt phenotypes of very different severity, from complete sterility in just a few generations (Spracklin et al., 2017), to little or no Mrt phenotype (this study). Additionally, *nrde-1* and *nrde-4* mutants become progressively sterile when maintained at 20°C whilst the phenotype in *nrde-2* and *hrde-1* mutants only becomes apparent at 25°C (Buckley et al., 2012). These variations do not appear to correlate with the severity of the TEI defect. Further research is required to explain these differences and determine the precise roles for the TEI genes in germline maintenance.

It has been suggested that a decreased cost of germline maintenance may cause lifespan extension through the reallocation of resources from germline maintenance to somatic maintenance (Maklakov and Immler, 2016). Our results are consistent with this model; we have shown that *set-32* mutants exhibit a Mrt phenotype implying decreased germline maintenance activity, and extended lifespan. Additionally, *set-25* mutants have apparently normal germline maintenance and normal lifespan. It would be interesting to investigate the lifespan of other TEI mutants displaying Mrt phenotypes to determine whether they also show lifespan extension, and hence whether the germline maintenance activity of the TEI genes has a general link to lifespan.

## Conclusion

We have shown that the chromatin modifiers SET-25 and SET-32 are required only to establish a heritable silencing signal and are dispensable for the maintenance of silencing across subsequent generations, implying a chromatin-independent mechanism maintains heritable silencing. Whilst 2° siRNA-associated factors including HRDE-1 and NRDE-2 are required for this maintenance, we have shown that the abundance of 2° siRNAs does not correlate with heritable silencing potential. This opens the field to the search for additional mechanisms responsible for the maintenance of transgenerational epigenetic inheritance.

## Acknowledgments

R.W. was supported by an Australian Government Research Training Program Scholarship, and A.A. by a Discovery Early Career Researcher Award. We thank the Australian Microscopy & Microanalysis Research Facility at the Australian Centre for Microscopy & Microanalysis (University of Sydney) for access to microscopes and assistance with imaging and analysis, and the Sydney Informatics Hub (University of Sydney) for providing access to High Performance Computing networks. We also thank the CGC (University of Minnesota) for strains, which is funded by NIH Office of Research Infrastructure Programs (P40 OD010440).

## Author Contributions

R.W. and A.A. conceived and designed the study and wrote the manuscript. R.W., G.B., M.H., D.H., M.L., P.R., and A.A. performed the experiments. R.W., M.L., and A.A. analysed the data. M.L., P.R., and A.A. provided expertise and feedback.

## Methods

### Strain list

Wild-type Bristol N2, SX461 *mjIs31[ppie-1::gfp::h2b]* II, AKA33 *set-32(smb11)* I; *mjIs31[ppie-1::gfp::h2b]* II, AKA35 *mjIs31[ppie-1::gfp::h2b]* II; *set-25(n5021)* III (outcrossed 6x from MT17463), AKA36 *set-32(ok1457)* I; *mjIs31[ppie-1::gfp::h2b]* II (outcrossed 6x from VC967), AKA37 *set-32(ok1457)* I; *mjIs31[ppie-1::gfp::h2b]* II; *set-25(n5021)* III, SX1442 *mjIs31[ppie-1::gfp::h2b]* II; *nrde-2(mj168)* II, SX2127 *mjIs31[ppie-1::gfp::h2b]* II; *hrde-1(tm1200)* III, AKA48 *mjIs31[ppie-1::gfp::h2b]* II; *mCherry::set-25+loxP(smb16)* III, PHX229 *set-32(syb229)* I, AKA59 *mjIs31[ppie-1::gfp::h2b]* II; *rhIs4[glr-1::gfp]* III; *him-8(e1489)* IV, AKA60 *set-32(ok1457)* I; *mjIs31[ppie-1::gfp::h2b]* II, *rhIs4[glr-1::gfp]* III; *him-8(e1489)* IV.

In all experiments strains were in *mjIs31[ppie-1::gfp::h2b]* background and SX461 was used as the wild-type, except in expression experiments (Figure 3A-D and Supplementary Figure S3A) where N2 was used as the wild-type.

### Cultivation and maintenance of *C. elegans*

Animals were cultured according to standard procedures (Brenner, 1974). Unless otherwise indicated, animals were grown on Nematode Growth Medium (NGM) (2% (w/v) agar, 50 mM NaCl, 0.25% (w/v) peptone, 1 mM CaCl_2_, 5 μg/ml cholesterol, 25 mM K_3_PO_4_ and 1 mM MgSO_4_ in H_2_O) plates seeded with OP50 *E. coli* bacteria, and experiments were performed at 20°C.

### *C. elegans* synchronisation

To obtain synchronised adults for extraction of small RNAs, gravid adults were bleached and the resulting staged embryos were plated and grown for 4 days. To obtain synchronised animals for other experiments, young adults laid embryos for 2 hours before being removed from plates. Resulting staged embryos were grown for ~60 hours to produce larval stage 4 (L4) animals.

### CRISPR/Cas9

All plasmid sequences were confirmed by Sanger sequencing and purified with DNA Clean and Concentrator-5 (Zymo Research). sgRNA target sequences were designed and incorporated into a *pU6::klp-12* sgRNA expression vector by PCR as previously described (Friedland et al., 2013; Norris et al., 2015). To create the *set-32(smb11)* deletion, an injection mix was injected into gonads of young adult animals consisting of sgRNA expression vector (150 ng/μL), Cas9 expression vector (*peft-3::cas9::tbb-2*) (150 ng/μL) (a kind gift from the de Bono Lab), pCFJ90 (*pmyo-2::mCherry::unc-54*) (5 ng/μL) and pCFJ104 (*pmyo-3::mCherry::unc-54*) (5 ng/μL). PCR was performed on the genomic DNA of mCherry positive animals to identify deletions. To create the *mCherry::set-25* strain (AKA48) we used the strategy outlined in Norris et al. (2015). Briefly, a repair template was constructed from approximately 950 base pair homology arms cloned into a loxP_myo2_neoR_mCherry_intron repair template vector also containing a neomycin resistance gene and *pmyo-2::GFP* (Norris et al., 2015). Homology arms contained synonymous mutations at the sgRNA target site. An injection mix was injected into gonads of young adult animals consisting of the sgRNA expression vector (200 ng/μL), repair template (70 ng/μL), Cas9 expression vector (45 ng/μL), pCFJ90 (5 ng/μL) and pCFJ104 (7 ng/μL). Transgenic animals were identified by survival after addition of G418 (500 μL of 25 mg/mL solution added to standard 6 cm plates). Once integrated lines were confirmed, animals were injected with pDD104 (*peft-3::Cre*) (50 ng/μL) and pCFJ90 (5 ng/μL). Successful Cre recombination was detected by the loss of pharyngeal GFP. Correct integration and Cre recombination was confirmed by Sanger sequencing across the homology arms. The *set-32::mCherry* strain was generated by SunyBiotech (China) and correct integration confirmed by sequencing.

### RNAi inheritance assays

RNAi was performed by feeding as previously described (Kamath et al., 2001). HT115(DE3) bacteria carrying IPTG-inducible L4440 (empty vector) or L4440-*gfp* plasmids was grown at 37°C for 7-8 hours with 100 μg/mL ampicillin. Cultures were seeded on NGM plates containing 25 μg/mL carbenicillin and 1 mM IPTG grown overnight at room temperature. Young adults were plated onto RNAi bacteria and their progeny scored for the presence of GFP as adults 4 days later. Silenced adults were transferred to OP50 plates to produce subsequent generations.

To test for a requirement in the P0 or F1 generation, *set-32(ok1457)* or *set-25(n5021)* L4 hermaphrodites were crossed to wild-type males on OP50 plates. 24 hours later, fertilised adult hermaphrodites were plated onto RNAi food plates prepared as above. 3 days later, L4 hermaphrodite progeny (the P0 generation) were moved to new RNAi plates to prevent mating with male progeny. 24 hours later, adult hermaphrodites were transferred to OP50 plates to produce the F1 generation. The P0 adults were then lysed and genotyped to confirm heterozygosity, and GFP-positive F1 offspring were lysed and genotyped.

Each experiment was performed in triplicate (5 independent plates per replicate), with ~100 animals per generation scored in each replicate. Scoring was performed blind to the genotype of the strains.

### Small RNA sequencing

The RNAi inheritance assay was performed as above, with the exception that populations were created by plating bleached embryos. RNAi-treated P0 animals (~200 per replicate), and F1 animals (~20 per replicate) sorted into GFP-expressing or ‐silenced groups, were collected in 1 mL TriSure (Bioline). Animals were cracked by five freeze/thaw cycles in liquid nitrogen, and RNA extracted using the manufacturer’s instructions with the exception that 1 uL Glycogen (20 mg/mL, Roche) was added as a carrier during the precipitation step, which was performed overnight at 20 °C. 250 – 500 ng of RNA was treated with 5′ polyphosphatase (Epicentre) following the manufacturer’s instructions after which small RNA libraries were prepared using the NEBNext Multiplex Small RNA Library Prep Set essentially as described in the manual. Sequencing of libraries was performed by AGRF on an Illumina HiSeq2500. The experiment was performed in duplicate except for strains AKA33 and SX2127 for which the P0 generation was performed in triplicate.

### Bioinformatic analysis

All initial QC, trimming and mapping analysis was performed using CLC Genomics (Qiagen). Libraries were trimmed to remove adapters and filtered for a quality score of >30. Reads were mapped to WormBase release WS260 and the GFP transgene with a maximum of one mismatch. For the GFP-mapping siRNA analysis, BAM files were exported from CLC Genomics and analysed in R using a combination of the viRome package (Watson et al., 2013) and custom scripts. For endo siRNA analysis, the number of reads mapping to particular biotypes was determined in CLC Genomics using WormBase WS260 annotations and extracted. Data analysis and visualisation was performed using Excel and GraphPad Prism. Sequence data has been submitted to the SRA at NCBI (BioProject ID PRJNA431114).

### Microscopy and analysis

Animals were scored for the presence or absence of GFP with a Nikon SMZ18 Microscope.

For Figure 3A-E and Figure S3A, animals were immobilised in M9 with 0.2% Tetramizol and mounted on glass slides. DIC and fluorescent imaging was performed using standard methods using a Leica Sp5 Multiphoton confocal microscope (SET-32::mCherry and mCherry::SET-25) and a Nikon Ti-E spinning disk microscope (mCherry::SET-25). For SET-32::mCherry analysis, all images of the tagged strain and N2 control animals were taken under identical conditions. Quantification was performed using ImageJ and statistical tests performed in Excel.

For immunofluorescence experiments in Figure 3F-H and Figure S3B-D, germlines were imaged using a Nikon C2 Basic Confocal microscope. Within conditions (DAPI, H3K9me2, and H3K9me3 staining) all strains were imaged under identical parameters. Average fluorescence intensity was quantified using ImageJ (n=25-39). An intensity threshold-based mask for nuclei was generated using the DAPI signal then used to determine the average intensity of H3K9me3 and H3K9me2 staining in nuclei, to control for variable space between nuclei in different germlines. In all strains we observed lower average intensity of H3K9me3 in the mitotic zone of the germline compared with the pachytene zone and so quantified intensity for each separately: average intensity was measured for the mitotic and pachytene zones of the germline separately by drawing a region around the whole mitotic zone, and around the beginning of the pachytene zone of equal size (by number of cells) to the mitotic region (n=24-39). The ratio of intensity in the mitotic:pachytene zones was calculated by dividing the mitotic intensity by the pachytene intensity for each germline. H3K9me2 staining was consistent throughout the germline, so the mitotic region was used for quantification. Intensity in individual nuclei was quantified by drawing individual regions around the 20 most distal germline nuclei in a single plane in 4 representative animals per strain.

For Figure 4E-G, synchronised L4 hermaphrodites were incubated for 12 or 24 hours, then fixed as previously described (Nishimura et al., 2015). DNA was visualised with DAPI. Imaging was performed using an Olympus FluoView FV1000 Confocal microscope, and processed using Olympus FluoView software. A Z-stack of images from one spermatheca per hermaphrodite was collected and flattened to create a maximum intensity projection image using ImageJ. Sperm from one spermatheca per hermaphrodite were counted as previously described (Nishimura et al., 2015) (n=6 for 12 hours, n=10-11 for 24 hours).

### Germline immunofluorescence

One-day-old adult hermaphrodites were dissected in M9 with 0.05% Tetramizol to release gonads onto poly-L-lysine coated slides. Germlines were cracked by freeze/thawing in liquid nitrogen, then fixed in −20°C methanol for 1 min followed by 3.7% paraformaldehyde, 1xPBS, 0.08 M HEPES (pH 6.9), 1.6 mM MgSO_4_, 0.8 mM EGTA for 30 min. Primary antibodies used were rabbit polyclonal to Histone H3 (tri methyl K9) (Ab8898, Abcam) and mouse monoclonal to Histone H3 (di methyl K9) (Ab1220, Abcam), diluted 1:300 in 30% normal goat serum in PBS. Secondary antibodies used were goat anti-rabbit Alexa Fluor 488 and rabbit anti-mouse Alexa Fluor 555 (Invitrogen), diluted 1:1000 in 30% normal goat serum in PBS. DNA was visualised with DAPI.

### Protein Subcellular Fractionation

Large-scale populations of animals were grown on enriched peptone plates (20 mM NaCl, 20 g/L peptone, 25 g/L agar, 5 μg/mL cholesterol, 1 mM MgSO_4_, 25 mM K_3_PO_4_) seeded with NA22 *E. coli* bacteria. Animals were washed several times in M9 buffer before homogenisation. Approximately 20,000 whole worms per strain were fractionated using a detergent solubility-based kit designed for tissue separations (Pierce Tissue Subcellular Fractionation Kit, Thermo). Briefly, whole worms were resuspended in 1 mL of cytosol extraction buffer containing protease inhibitors, combined with an equal volume of 0.7 mm zirconia beads in a 2 mL screw-cap tube, and bead-beated for 5 seconds at 4oC using in a BioSpec Products MiniBeadBeater-24. This extract was fractionated according to manufacturer’s instructions for the Pierce Tissue Subcellular Fractionation Kit (Thermo). The protein content in each fraction was quantified by a BCA protein assay (Thermo).

### Protein digestion, peptide clean-up and quantitation

Proteins from the chromatin fraction (50 ug) were denatured and reduced in 2% SDS and 10 mM triscarboxyethylphosphine (TCEP), and 20 mM sodium phosphate buffer pH 6.0 and 150 mM NaCl. The protein samples were heated to 65oC in a ThermoMixer-C (Eppendorf) for 10 min at 1000 rpm. Once cooled to room temperature, N-ethylmaleimide (NEM) was added to the fractions at a final concentration of 20 mM and allowed to incubate for 30 min at room temperature. The fractions were buffer exchanged and trypsin digested using the SP3 method described previously (Hughes et al., 2014).

### LC-MS/MS and analysis of spectra

Using a Thermo Fisher Scientific EasyLC 1200 UHPLC, peptides in 4% (vol/vol) formic acid (injection volume 3 μL, approximately 500 ng peptides) were directly injected onto a 50 cm × 75 μm reverse phase C18 column with 1.9 μm particles (Dr. Maisch GmbH) with integrated emitter. Peptides were separated over a gradient from 4% acetonitrile to 32% acetonitrile over 30 min with a flow rate of 300 nL min^−1^. The peptides were ionized by electrospray ionization at +2.3 kV. Tandem mass spectrometry analysis was carried out on a Q-Exactive HF mass spectrometer (Thermo Fisher Scientific) using HCD fragmentation. The data-dependent acquisition method used acquired MS/MS spectra on the top 5 most abundant ions at any one point during the gradient. All the RAW MS data have been deposited to the ProteomeXchange Consortium (http://proteomecentral.proteomexchange.org) via the PRIDE partner repository with the dataset identifier PXD008754, username, password: 9GD5YNDU. The RAW data produced by the mass spectrometer were analysed using Proteome Discoverer 2.2 (Thermo) and the Byonic Search Engine (Protein Metrics). Peptide and protein level identification were both set to a false discovery rate of 1% using a target-decoy based strategy. The database supplied to the search engine for peptide identifications was a combined *C. elegans* and *E. coli* Swissprot database downloaded on the 11th April 2017. The mass tolerance was set to 3 ppm for precursor ions and MS/MS mass tolerance was set at 10 ppm. Enzyme was set to trypsin (cleavage C-terminal to R/K) with up to 3 missed cleavages. Deamidation of N/Q, oxidation of M were set as common variable modifications of which only 1 was allowed. N-terminal pyro-E/Q, protein N-terminal acetylation, acetylation of K, methylation of K/R, dimethylation of K/R, trimethylation of K were set as rare variable modifications of which only 2 were allowed. N-ethylmaleimide on C was searched as a fixed modification. The output from the Byonic search has also been uploaded to the ProteomeXchange Consortium under the same identifier given above.

### Histone Peptide Quantitation

The ratio of each modified peptide to a control peptide (either the cognate unmodified peptide, or an unmodified peptide from elsewhere in the protein) was calculated from extracted ion chromatograms of each, across all samples. The area under each peak was integrated and ratios calculated. Significance was calculated using two-way ANOVA in GraphPad Prism.

### Mortal germline assays

Animals were maintained at 20°C for at least 5 generations before being shifted to 25°C at L3 stage. 20 replicate lines were created from individual animals and maintained as separate populations throughout the experiment. A single L3 animal per line was picked to a new plate to create each subsequent generation. In the regular mortal germline assay, the mean brood size per strain was calculated at each generation by averaging the number of progeny between lines. When a sterile individual arose resulting in no progeny to produce the next generation, the line was discarded. For the purpose of data analysis discarded lines were recorded as having zero progeny for all subsequent generations. In the cumulative sterility assay, the number of times sterility arose was counted. When a sterile individual arose, that individual was replaced with an animal from a backup population which had been maintained at 25°C for the same number of generations in order to retain a consistent number of lines. For both assays, scoring was performed blind to the strain genotype.

### Brood size assays

To measure regular brood size, L4 hermaphrodites were plated onto growth plates and transferred every 12 hours for the first three days, then every 24 hours until they had stopped laying or died. Plates from which animals had been transferred were incubated for 48 hours, then the numbers of live progeny and unfertilised eggs scored.

To test for a male germline defect, *set-32(ok1457)* or wild-type control L4 hermaphrodites were mated to wild-type L4 males for 24 hours. Males were then removed, and hermaphrodites transferred every 12 hours for the first three days, then every 24 hours until they had stopped laying or died. Live progeny were scored as above. The percentage of male progeny was monitored, and only the progeny of successfully-mated hermaphrodites were included in analysis, indicated by the presence of ~50% males.

To test for a defect in male sperm, wild-type L4 hermaphrodites were mated to control *him-8(e1489)* L4 males or mutant *set-32(ok1457); him-8(e1489)* L4 males carrying an integrated *glr-1::gfp* neuronal reporter transgene for 24 hours. Males were then removed, and hermaphrodites transferred every 24 hours until they had stopped laying or died. Numbers of GFP-positive (cross-progeny) and ‐negative (self-progeny) live offspring were scored.

Each experiment was performed in duplicate with n=10 animals at Day 0 per replicate. Animals which died or were lost within the first 24 hours of adulthood were excluded from analysis. Scoring was performed blind to the strain genotype.

### Lifespan assays

Synchronised L4 animals were plated on growth plates in the absence (n=110) or presence (n=100) of FUDR (100 μM). In the absence of FUDR, animals were transferred to new plates every day for the first 8 days to separate adults from progeny, and then once per week until death. In the presence of FUDR, animals were transferred once after 10 days. Animals were scored daily and considered dead when they did not respond to gentle touch with a platinum wire. Animals that displayed vulval rupture or progeny hatching within the parent were removed from plates and censored from analysis. Scoring was performed blind to the strain genotype.

## Supplemental Information Titles and Legends

**Figure S1. The *set-25* and *set-32* loci and mutant alleles. A.** Schematic representing the *set-25* and *set-32* transcripts. Exons are represented by black boxes, introns by connecting lines, and untranslated regions by white boxes. The sequence encoding the SET domain is indicated in yellow. Brackets indicate deleted sequence in mutant alleles. **B.** Predicted protein for *set-25*, *set-32* and mutants. The location of the SET domain is indicated in yellow. aa denotes length in amino acids.

**Figure S2. Extended small RNA analysis. A.** Small RNAs (reads per million) in the wild-type strain that map to the indicated biotype. **B-D.** Small RNAs (reads per million) with the indicated biotype. Strains are indicated: wt is wild-type, *s25* is *set-25(n5021)*, *s32^CR^* is *set-32(smb11)*, *s32^Del^* is *set-32(ok1457)*, *nrde* is *nrde-2(mj168)*, *hrde* is *hrde-1(tm1200)*. The top panel shows P0 control and RNAi-treated combined (P0) and F1 GFP-expressing and GFP-silenced animals combined (F1). Two-way ANOVA was performed and results for variance at the genotype or generation level is shown. The bottom panel shows P0 and F1 individually, normalised to the relevant control (dashed black line). Comparisons were performed by two-way ANOVA with Tukey’s post hoc. * p<0.05 ** p<0.01 **** p<0.0001

**Figure S3. SET-32 expression patterns and H3K9 methylation analysis. A.** The entire germline of representative SET-32::mCherry expressing animals and wild-type animals of the indicated age. Germline is outlined in white. **B.** Representative single confocal plane images of the mitotic zone of dissected adult gonads of the indicated strain. Panels show anti-H3K9me2 staining (yellow), DAPI staining (blue), and an overlay of H3K9me2 and DAPI staining, as indicated. Strains were imaged under identical parameters within staining conditions. Scale bars in **A-B** represent 20 μm. **C.** Quantification of fluorescence intensity of H3K9me2 staining in the mitotic zone. ImageJ was used to generate an intensity threshold-based mask for nuclei using the DAPI signal, then the average intensity of H3K9me2 staining in nuclei was measured. Data are mean ±SEM, n=24-29. Comparisons were performed by one-way ANOVA with Tukey’s post hoc. **** p<0.0001 **D.** Quantification of fluorescence intensity of H3K9me3 staining in individual nuclei. ImageJ was used to generate an intensity threshold-based mask for nuclei using the DAPI signal, then the average intensity of H3K9me3 staining in individual nuclei was measured. Data are mean ±SEM, n=80 nuclei from 4 germlines. **E.** Quantification of H3K79 levels in the indicated mutant strains by mass spectrometry. For each modification, levels were normalised to a relevant control peptide and are displayed as % of wild-type (indicated by dotted line). Data are mean ±SEM, n=3 replicates for each mutant strain. Comparisons were performed by two-way ANOVA with Dunnett’s post hoc. * p<0.05

**Figure S4. *set-32* mutants display a mortal germline phenotype, defective sperm, and extended lifespan. A.** The number of times a line went sterile during maintenance at 25°C was counted, and the sterile line replaced by an animal from a backup population. Data are mean ±SEM, n=20 lines. **B-C.** The average number of **B.** live progeny and **C.** unfertilised eggs produced per animal per day. Data are mean ±SEM per day, n=18-20. **D.** Brood sizes of wild-type or *set-32(ok1457)* hermaphrodites mated to wild-type males. Data are mean ±SEM per day, n=19. **E.** Wild-type hermaphrodites were crossed to males of the indicated genotype carrying a neuronal GFP transgene. Cross-progeny (expressing the GFP transgene) and self-progeny (not expressing GFP) were counted. Data are mean ±SEM, n=20-26. Comparisons were performed by two-way ANOVA with Sidak’s post hoc. **** p<0.0001 **F.** Replicate of assay in Figure 4H; lifespan assay performed in the absence of FUDR, n=110 at Day 0. **G-I.** Independent replicate lifespan assays performed in the presence of FUDR, n=100 at Day 0. **J.** Median lifespan of each strain and p value of the survival curves of each mutant strain compared with wild-type. Comparisons were performed by Log-rank test. * p<0.05 ** p<0.01 *** p<0.001 **** p<0.0001

